# A novel toolbox of GATEWAY-compatible vectors for rapid functional gene analysis in soybean composite plants

**DOI:** 10.1101/2024.10.12.617978

**Authors:** Joffrey Mejias, Alexandra Margets, Melissa Bredow, Jessica Foster, Ekkachai Khwanbua, Jackson Goshon, Thomas R. Maier, Steven A. Whitham, Roger W. Innes, Thomas J. Baum

## Abstract

The generation of transgenic plants is essential for plant biology research to investigate plant physiology, pathogen interactions and gene function. However, producing stable transgenic plants for plants such as soybean is a laborious and time-consuming process, which can impede research progress. Composite plants consisting of wild-type shoots and transgenic roots are an alternative method for generating transgenic plant tissues that can facilitate functional analysis of genes-of-interest involved in root development or root-microbe interactions. In this report, we introduce a novel set of GATEWAY-compatible vectors that enable a wide range of molecular biology uses in roots of soybean composite plants. These vectors incorporate in-frame epitope fusions of green fluorescent protein, 3x-HA, or miniTurbo-ID, which can be easily fused to a gene-of-interest using the GATEWAY cloning system. Moreover, these vectors allow for the identification of transgenic roots using either mCherry fluorescence or the RUBY marker. We demonstrate the functionality of these vectors by expressing subcellular markers in soybean, providing evidence of their effectiveness in generating protein fusions in composite soybean plants. Furthermore, we show how these vectors can be used for gene function analysis by expressing the bacterial effector, AvrPphB in composite roots, enabling the identification of soybean targets via immunoprecipitation followed by mass spectrometry. Additionally, we demonstrate the successful expression of stable miniTurbo-ID fusion proteins in composite roots. Overall, this new set of vectors is a powerful tool that can be used to assess subcellular localization and perform gene function analyses in soybean roots without the need to generate stable transgenic plants.

**Key Message:** We developed a set of GATEWAY vectors to accelerate gene function analysis in soybean composite plants to rapidly screen transgenic roots and investigate subcellular localization, protein-protein interactions, and root-pathogen interactions.

## Background

Soybean (*Glycine max*) is the most widely grown oil seed crop worldwide and is a major source of food for humans and livestock (Pagano and Miransari 2016). Genetic transformation of soybean is less efficient and more time-consuming than many other crop species, causing a major bottleneck for high-throughput gene function analysis (Xu et al. 2022). Transgenic expression of genes using *in vitro* hairy roots has been widely used as an alternative to stable transgenics (Chen et al. 2018); however the generated roots are grown on media instead of in soil and are multiplied without aerial tissues, which is labor-intensive and could introduce bias when assessing a phenotype on a given mutated gene. More recently, soybean composite plants have been demonstrated as an efficient alternative for generating transgenic roots and performing functional analysis of genes (Fan et al. 2020; Matthews and Youssef 2016). Composite plants can be generated rapidly without the need for aseptic growth conditions employed during generation of hairy roots, offering a more efficient method for obtaining transgenic roots that still remain attached to wild-type aerial tissues (Fan et al. 2020).

Cloning a gene-of-interest (GOI) using the classical restriction digestion-ligation method can require multiple cloning steps, especially when the GOI requires fusion to a particular epitope. GATEWAY technology is a method of choice to rapidly clone and express a gene *in frame* with a given epitope when a set of destination vectors are available (Curtis and Grossniklaus 2003). In plants, different sets of GATEWAY-compatible destination vectors are available that harbor expression cassettes or reporter genes used for multiple purposes. The most widely used set of vectors for *in planta* expression are those developed by (Nakagawa et al. 2007) and (Karimi, Inzé, and Depicker 2002), which drive gene expression under a viral P35S constitutive promoter. These vectors contain the *aadA* resistance cassette, a gene encoding for an adenylyltransferase that confers bacterial resistance to spectinomycin and streptomycin. This system is compatible with the widely used *Agrobacterium tumefaciens* strain GV3101, allowing for the transient expression of chimeric proteins in *Nicotiana benthamiana* and the generation of stable *Arabidopsis thaliana* transgenic lines.

Our aim was to adapt the GATEWAY cloning system, which allows for the efficient shuttling of a GOI in a single step reaction into a given plant destination vector, for the purposes of generating soybean composite plants in a high throughput gene function analysis system. Soybean composite plants are produced using the virulent *A. rhizogenes* strain K599 (Mankin et al. 2007) which has high resistance to both spectinomycin and streptomycin encoded on a helper vector, and also has cryptic resistance to gentamicin. Importantly, the most commonly used entry GATEWAY vectors, pDONRs (pDONR207 and pDONR221; Thermo Fisher Scientific), carry resistance to gentamicin and kanamycin, respectively, rendering the set of vectors developed by Nakagawa et al. (2007) and Karimi et al. (2002) incompatible with the use of GATEWAY cloning system in the *A. rhizogenes* K599 strain. Currently, the only available GATEWAY vectors compatible with K599 strain are the pRAP (Klink et al. 2021) and pJan (Alzohairy, MacDonald, and Matthews 2013) vector series which harbor tetracycline resistance. However, they do not contain any pre-assembled *in-frame* epitope tags (only a AttR1-ccdB-AttR2 cassette). Moreover, root selection is performed based on green fluorescence protein (GFP), requiring that each root system is screened under an epifluorescence microscope.

We, therefore, generated a novel set of GATEWAY-compatible vectors carrying a tetracycline resistance gene for the generation of composite soybean plants using the *A. rhizogenes* strain K599. In the GATEWAY cloning system, the ccdB cassette encodes a toxin that kills *Escherichia coli* strains that do not possess the corresponding antitoxin gene (Van Melderen 2002) as well as a chloramphenicol resistance cassette. This ccdB cassette is widely used in the GATEWAY cloning system to counter-select bacteria that did not receive the GOI. Our novel set of GATEWAY destination vectors includes different in-frame epitopes with the ccdB recombination cassette allowing GOI expression, analysis of subcellular localization, or identification of protein interactors through proximity-labeling or immunoprecipitation (IP) approaches. Along with the GOI expression constructs, we incorporated on the same T-DNA the fluorescent reporter gene mCherry as well as the RUBY reporter gene cassette (He et al. 2020), simplifying the identification of transgenic roots. A strong constitutive promoter from soybean, *Gm*Ubi (Ubiquitin) (De La Torre and Finer 2015), drives the expression of the GOI in the current vectors but can be easily replaced with another promoter using GC-rich restriction enzymes. Expression of the mCherry and RUBY markers is driven by the cassava vein mosaic virus promoter CsVMV which displays high constitutive expression in soybean roots (Govindarajulu et al. 2008). We also generated a similar set of vectors where the *Gm*Ubi promoter was replaced by a dexamethasone (DEX)-inducible promoter (Vinatzer et al. 2006) to fine-tune expression of a GOI, which could be used to drive the expression of toxic genes.

## Materials and Methods

### Vector construction

The pICH75322_Kan^R^ vector (Addgene #48051) was digested with *SspI* and *AfeI* restriction enzymes to remove the kanamycin resistance cassette. The tetracycline resistance cassette was amplified from the pBR322 vector (Promega), and *SspI* restriction sites were added to both sides of the PCR amplicon which was subsequently ligated into the digested vector and transformed into *E. coli* DH5α cells. The resulting vector was confirmed by Sanger sequencing and named pICH75322_Tet^R^. The following Golden Gate (GG) modules were amplified from the pG2RNAi2 vector (Noon et al. 2016): *Gm*Ubi promoter (p*Gm*Ubi), the Rubisco terminator (Trbsc) and the CsVMV promoter (pCsVMV). The enhanced GFP (eGFP) and nopaline synthase terminator (TNos) modules were amplified from the pK7WGF2 vector (Karimi, Inzé, and Depicker 2002). The mCherry module was amplified from the pENTR221-mCherry vector (Addgene #79509), and the RUBY module was amplified from the P35S:RUBY vector (Addgene #160908). For each module, compatible *BsaI* restriction sites were added through PCR amplification, as well as extra restriction enzyme sites for the *Gm*Ubi and eGFP modules, and amplicons were independently cloned in the TA-cloning vector pMiniT2.0 (*BsaI* free) following the manufacturer’s instruction (New England Biolab, Massachusetts, USA). Inserted sequences were confirmed by Sanger sequencing at the Iowa State University (ISU) DNA facility using the Mini-F and Mini-R primers (Table S1). Compatible GG modules were used (75 ng each), as well as the destination vector pICH75322_Tet^R^ (75ng), in a GG reaction for 35 cycles to generate the two intermediate vectors: p*Gm*Ubi:eGFP-Trbsc + pCsVMV:mCherry-TNos and p*Gm*Ubi:GFP-Trbsc + pCsVMV:RUBY-TNos. To convert these two intermediate vectors into GATEWAY-compatible vectors, the eGFP module was removed by digestion using *KpnI* and *AvrII*. The eGFP-ccdB, ccdB-eGFP, 3xHA-ccdB, ccdB-3xHA and ccdB modules were amplified from pK7GWF2, pK7WGF2, pGWB414, pGWB415 and pK2GW7, respectively (Karimi, Inzé, and Depicker 2002); (Nakagawa et al. 2007) and *KpnI* and *AvrII* restriction sites were added by PCR amplification. Each ccdB cassette was inserted in the digested intermediate vector by ligation, and each vector was sequenced using whole plasmid sequencing (Plasmidsaurus, Oregon, USA). To construct the vectors containing the miniTurboID-V5-ccdB and ccdB-miniTurboID-V5 cassette, we created two intermediate vectors using a similar GG strategy where the GFP module was replaced by the miniTurboID-V5 fused to a linker (GGGS) for which we added consecutive *KpnI* and *AvrII* restriction sides sites upstream or downstream depending on the orientation of the cassette. The miniTurboID gene was amplified from the pBSDONR (P4r-P2):miniTurboID-V5 vector (Margets et al. 2024). We then digested these intermediate vectors using *KpnI* and *AvrII* and inserted the ccdB cassette using restriction-ligation.

For promoter analysis using RUBY as a screenable marker, we generated a p*Gm*Ubi:RUBY-Trbsc + pCsVMV:mCherry-TNos vector using similar GG strategy as described previously where the eGFP module was replaced by a RUBY module. Subsequently, we digested this vector using *ApaI* and *AscI* to remove the *Gm*Ubi promoter and amplified p*Gm*Actin from soybean gDNA. The P35S+Ωenhancer was PCR amplified from the pDOE-5 vector (Gookin and Assmann 2014), and *ApaI* and *AscI* restriction sites were added using specific primer and inserted using restriction-ligation. To construct the p*Gm*NHL1:GUS-Trbsc + pCsVMV:mCherry-TNos, vector we used p*Gm*Ubi:ccdB-Trbsc + pCsVMV:mCherry-TNos as a destination vector in an LR reaction with a pDONR221 vector containing the GUS plus gene. The p*Gm*Ubi promoter was then replaced by the p*Gm*NHL1 promoter (1399 bp upsteam of ATG) which was amplified from soybean gDNA using restriction-ligation using the *ApaI* and *AscI* restriction sites.

To generate the subcellular localization marker vectors for microtubules (GFP-MAP4/MBD), actin filaments (LifeActin-mCherry), endoplasmic reticulum (AtWAK2-mCherry-HDEL), plasma membrane (PIP2-GFP), and plasmodesmata (PDLP1-GFP), previously described GATEWAY destination vectors (Ivanov and Harrison 2014) were used as templates for BP reactions. A back-BP reaction was performed to obtain all the pDONR221 entry vectors (pDONR221:MAP4/MBD; pDONR221:AtWAK2-mCherry-HDEL; pDONR221:LifeActin-mCherry; pDONR221:PIP2 and pDONR221:PDLP1) for these markers, except for NLS-YFP that was synthesized (TwistBioscience, California, USA). Subsequently, these markers were shuttled from the pDONRs into the corresponding destination vectors (pDONR221:MAP4/MBD into eGFP-ccdB + mCherry; pDONR221:AtWAK2-mCherry-HDEL into ccdB + RUBY; pDONR221:LifeActin-mCherry into ccdB + RUBY; pDONR221:PIP2 into ccdB-eGFP + mCherry and pDONR221:PDLP1 into ccdB-eGFP + mCherry) using LR reactions to generate *in-frame* fusions of the subcellular markers with fluorescent tags.

To generate the eGFP-3xHA and the AvrPphB^C98S^-3xHA modules used for the IP experiments, the coding sequences of eGFP and AvrPphB^C98S^, without stop codons, were cloned into pDONR221 followed by an LR reaction into the destination vector p*Gm*Ubi:ccdB-3xHA + pCsVMV:RUBY.

To generate DEX-inducible vectors, the same approach as described for the first set of destination vectors was employed, using a DEX promoter synthesized without the endogenous *BsaI* and *AvrII* restriction sites (Eurogentec, California, USA). The synthesized sequence was adapted from the pTA7001 vector (Addgene #71745), but the p*Gm*Ubi promoter was used to drive the expression of the GVG fusion instead of P35S (p*Gm*Ubi:Gal4BD-VP16-GVG-TPeaRbsc-6xUAS-minimal P35S (P35Smin)). The complete DEX promoter was used in GoldenGate reactions to generate two intermediate vectors that were subsequently converted into GATEWAY vectors by restriction-ligation using *KpnI* and *AvrII* as mentioned above.

### Composite plant generation and maintenance

Soybeans plants were grown in a controlled chamber at 24°C at 45-65% relative humidity and a 16h/8h light/dark cycle. Williams 82 (W82) soybean seeds were sterilized using chlorine gas as described previously (Narayanan et al. 1999) and placed in pre-wet soil at 0.5 cm depth and covered for a week in a growth chamber. Seven days after sowing, trays with pots were prepared by placing a wet paper towel at the bottom of each pot before filling with sterilized vermiculite to avoid leakage. A hole was made at the center of each pot and 5 mL of K599 liquid culture, previously centrifuged and diluted to an O.D_600_ = 1.0 in 0.25X Gamborg’s medium (pH 5.75), was pipetted into each hole. Using a sterile razor blade, each soybean seedling was cut diagonally directly below the two cotyledons, and a paste of *Agrobacterium* culture (from plates) was applied to the wound site. The seedlings were then placed in the vermiculite trays which were filled with 1-2 cm of 0.25X Gamborg’s (pH 5.75) and covered with a lid for two weeks to ensure high humidity which promotes root generation. After two weeks, the plants were removed from the vermiculite, and the roots were monitored for mCherry fluorescence under an epifluorescence binocular microscope or assessed visually for RUBY production, and the non-transgenic roots were removed. The plants were immediately transplanted back into the vermiculite pots. The same process to remove non-transgenic roots was repeated one week later. A detailed schematic of soybean composite plant generation is shown in Figure 1.

**Figure 1.**
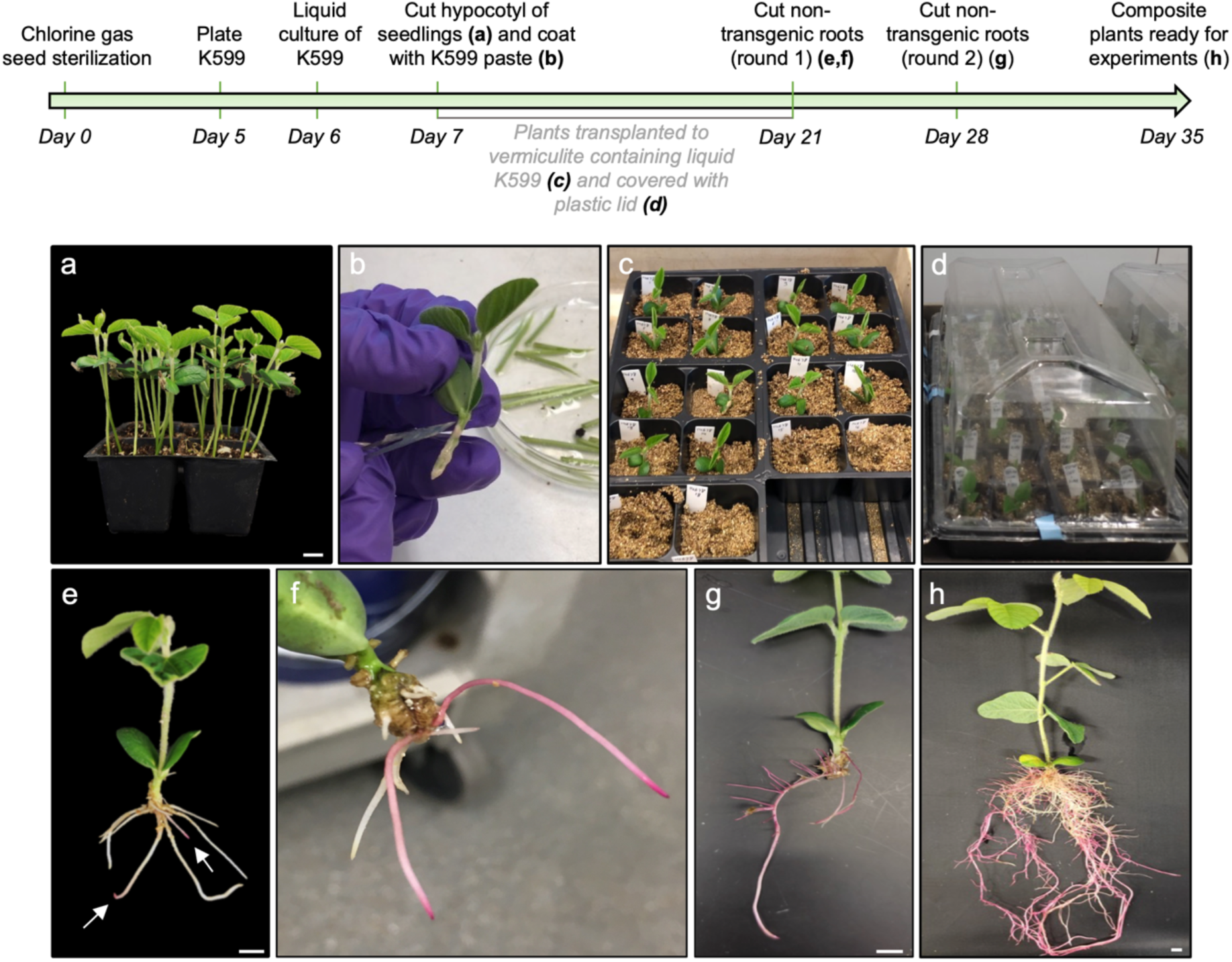
Soybean composite plant generation. Experimental timeline is shown at the top in green. (a) Soybean seedlings are grown for seven days. (b) A seedling is cut diagonally below the two cotyledons and a paste of Agrobacteria is applied to the wound site. (c) Soybean seedlings are transplanted into pre-made holes in vermiculite containing 5 mL of liquid K599 culture. (d) Plantlets are covered with a plastic lid for two weeks to maintain a high humidity environment. (e) Soybean plantet showing transgenic (pink) and non-transgenic (white) roots 21 days after germination. Transgenic roots are indicated with white arrows. (f) Close-up of composite plant showing two transgenic roots expressing the RUBY marker (pink) 21 days after germination. (g) Soybean composite plant after trimming non-transgenic roots 28 days after germination. (h) Soybean composite plant after two rounds of non-transgenic root trimming ready for experiments. Scale bar = 1 cm.

### Dexamethasone induction

Five weeks after inoculation with *A. rhizogenes* K599, soybean plants were removed from vermiculite and their roots were rinsed in water. The plants were then placed in a beaker filled with 200 mL of 0.25X Gamborg’s (pH 5.75) containing 50 µM of DEX. The roots were vacuum infiltrated for 1 min and then transferred to growth chambers. Medium containing DEX was refreshed every 24 h to minimize microbial growth. Roots were sampled at 6, 24, 48 and 96 h after induction, as indicated in figures.

### Confocal microscopy

The root tissues of composite soybeans and 14 day-post inoculation (dpi) galls induced by root-knot nematodes (RKN) were collected by hand under an epifluorescence binocular microscope when mCherry was used to screen transgenic roots, and longitudinally cut and observed under a LSM700 (Zeiss) confocal microscope. For clearing, soybean roots were cut longitudinally using a razor blade and fixed in 1X phosphate buffered saline (PBS) solution containing 4% paraformaldehyde for 24 h, with gentle shaking. The roots were washed three times for 1 min in 1X PBS and incubated in ClearSee (Kurihara et al. 2015) for one week, with gentle rocking, and transferred to ClearSee + 0.01% Calcofluor White for 1 h. Roots were then rinsed for 30 min in ClearSee before observation under a LSM700 confocal microscope. For simultaneous GFP/mCherry/Calcofluor imaging, samples were excited at 488 nm for GFP, 543 nm for mCherry, and 405nm for Calcofluor White in the multi-track scanning mode. GFP, mCherry, and Calcofluor White were detected selectively with 505–530 nm, 560–615 nm, and 450-490 nm band-pass emission filters, respectively.

### Immunoblot analysis and immunoprecipitation

For protein extractions for IP experiments, 1.5-2 g of transgenic roots collected from composite plants were ground in a pre-chilled mortar with pestle under liquid nitrogen and transferred to a 15 mL falcon tube with 2X the volume of filter-sterilized co-immunoprecipitation (coIP) extraction buffer containing 10% glycerol, 25 mM Tris-HCl (pH 7.5), 1 mM EDTA, 150 mM NaCl, 10 mM dithiothreitol (DTT), one plant protease inhibitor cocktail tablet (Sigma-Aldrich, Massachusetts, USA), and 1 mM phenylmethylsulfonyl fluoride. For miniTurbo-V5 and eGFP detection, total protein was extracted from 1.5-2.0 g of liquid nitrogen ground transgenic root tissue using GTEN buffer (10% glycerol, 25 mM Tris-HCl (pH 7.8), 1 mM EDTA, 150 mM NaCl, 0.2% of polyvinylpyrrolidone (PVPP), 1 mM DTT, 0.05% NP40 and one plant protease inhibitor cocktail tablet (Sigma-Aldrich). Samples were rotated at 4°C for 30 min with 30 sec of vortexing every 10 min. Samples were centrifuged twice at 9,000 g for 10 min, at 4°C to collect soluble proteins. For IPs, composite roots expressing GFP:3xHA+RUBY or AvrPphB^C98S^:3xHA+RUBY were harvested approximately 5 weeks post transgenic root initiation for protein extraction in co-IP extraction buffer (10% glycerol, 25 mM Tris-HCl pH 7.5, 1 mM EDTA, 150mM NaCl, 200 (l DTT, 1% protease inhibitor cocktail, 1 mM PMSF). 1 mL of protein lysate was incubated with 30 µl of HA-trap agarose (Chromotek, Illinois, USA) equilibrated beads at 4°C, on a rotator, for 2 h. Beads were washed three times with a co-IP extraction buffer supplemented with 0.05% NP-40. Beads were transferred to a new 1.5 ml microcentrifuge tube and an additional wash was performed with the co-IP extraction buffer, without NP-40, to remove detergent to prevent interference with mass spectrometry analysis.

For immunoblot analysis bait proteins captured on HA-trap beads were boiled in 50 µL sodium dodecyl sulfate (SDS) loading buffer + 10% ꞵ-mercaptoethanol at 95°C for 10 min. Samples were centrifuged at 2,500 g for 5 min and protein extracts were analyzed by immunoblot analysis. Loading controls were obtained by imaging the stain-free polyacrylamide gel using the stain-free gel setting on a ChemiDoc Imaging System (Bio-Rad, California, USA). Proteins were transferred to nitrocellulose membranes (Cytiva, Massachusetts, USA) at 300 milliamps for 1 h. Membranes were blocked in 5% non-fat powdered dry milk for 1 h at room temperature on a shaker. Membranes were then incubated overnight, at 4°C, on a rocker with horseradish peroxidase (HRP)-conjugated HA antibody (Roche, Indiana, USA, Cat. No. 12 013 819 001). Membranes were washed three times in 1X Tris-Buffered Saline (TBS) with 0.1% Tween-20 (TBS-T). Membranes were then incubated with ProtoGlow chemiluminescent substrate (National Diagnostics, Georgia, USA) for 5 min and exposed on ChemiDoc Imaging System (Bio-Rad) using custom settings for exposure time. For immunoblotting of miniTurboID-V5 and miniTurboID-V5-eGFP, the same procedure was used as for the HA constructs except for the use of anti-V5 (Sigma Aldrich; Cat No. V8012) and monoclonal anti-GFP (Roche; Cat No. 11814460001) antibodies coupled with the use of an anti-mouse secondary antibody conjugated to HRP (Thermo Fisher Scientific; Cat No 62-6520).

### Histochemical localization using GUS

The histochemical staining of GUS enzyme activity was performed as described previously (Jefferson, Kavanagh, and Bevan 1987). Pictures of GUS-stained roots, as well as RUBY expressing-nodules, were taken using a Zeiss Stemi SV11 microscope with AxioCam HRc.

### Soybean cyst nematode (SCN) and RKN infection in composite plants

Six-week-old composite plants were transferred into 8-inch containers filled with a mixture of one-part sand to one-part field soil that had previously been steam sterilized. Two days later, each plant was infected with 2,000 stage-two juvenile SCN (TN10). Thirty days after infection, SCN cysts were collected as described previously (Kandoth et al. 2011). Pictures of cysts were taken using a Zeiss Stemi SV11 microscope with AxioCam HRc and Axiovision SE64 V4.9.1 software to produce images of 1040 x 1040 resolution on white background. The cysts were counted, and their size extracted using the Nemacounter software (Mejias et al. 2024). For RKN nematode infection assays, 500 juvenile stage-two worms were inoculated on soybean composite plants expressing the corresponding subcellular marker in containers with a mixture of one-part sand to one-part field soil that had previously been steam sterilized. Galls were hand collected under a binocular 14 days after infection.

## Results

### Generation of GATEWAY-compatible vectors for soybean composite plants

We built the set of GATEWAY compatible vectors by utilizing the GoldenGate vector pICH75322 as a backbone (Weber et al. 2011) (MoClo kit). This vector was selected due to: (i) its ability to replicate in *A. rhizogenes* K599; (ii) the presence of *BsaI* restriction sites suitable for GoldenGate reactions and (iii) its relatively small size compared to most other T-DNA vectors. Although the GreenGate vector series (Lampropoulos et al. 2013) is more compact than the pICH series, they require an additional helper vector in Agrobacteria for replication. For example, the widely used *A. tumefaciens* GV3101 strain needs to carry an additional pSOUP helper vector (tetracycline resistant) for the proper replication of pGreen vector series (Hellens et al. 2000), which is also absent in the *A. rhizogenes* K599 strain. Therefore, pICH75322, which contains the necessary components for plasmid replication in *A. rhizogenes* K599 strain, is one of the most compact vectors in this category.

The pICH75322_KanR in its current state could not be used in GATEWAY reactions since it carries a kanamycin resistance similar to the pDONR221 entry vector. We therefore modified the pICH75322 vector by replacing the kanamycin resistance with a tetracycline resistance cassette, enabling the selection of the K599 strain on Tet^5^. Using pICH75322_TetR as a backbone, we developed two intermediate vectors, each containing two transcriptional units on the same T-DNA through the GoldenGate cloning system. The first transcriptional unit contains eGFP driven by the constitutive *Gm*Ubi promoter, followed by the transcriptional terminator Rbsc. The second transcriptional unit contains the viral promoter pCsVMV that drives the expression of a reporter gene used for selection of transgenic roots (mCherry or RUBY) followed by the downstream transcriptional terminator TNos (Figure 2a). Using mCherry as a reporter gene to select transgenic roots requires a binocular microscope equipped with epifluorescence and the requisite filters. On the other hand, the RUBY gene is a reporter gene encoding for three enzymes each separated by a P2A proteolytic cleavage site sequence allowing the release of the three individual enzymes required to catalyze the conversion of tyrosine into betalain. The accumulation of betalain renders plant tissue red *in vivo* without the need of a fixation step or the addition of a substrate (He et al. 2020). Using RUBY as a reporter gene for transgenic root selection is, therefore, more time-efficient and avoids false positive selection that can occur when using a fluorescent reporter due to plant autofluorescence. Each module used to build the two intermediate vectors was pre-cloned into the TA-cloning vector BsaI-free pMiniT2.0. GC-rich restriction sites surrounding the *Gm*Ubi promoter in these intermediate vectors were added to allow easy replacement with any promoter of interest. Since most promoters are AT-rich, these GC-rich restriction sites are unlikely to be found in a promoter sequence of interest. The functionality of the two intermediate vectors p*Gm*Ubi:GFP:TerRbsc + pCsVMV:mCherry:TNos and p*Gm*Ubi:GFP:TerRbsc + pCsVMV:RUBY:TNos were assessed by transforming *A. rhizogenes K599* and producing soybean composite plants. We confirmed expression of the reporter genes using an epifluorescence microscope equipped with a GFP or mCherry filter or using brightfield microscopy for RUBY (Figure 2a).

**Figure 2.**
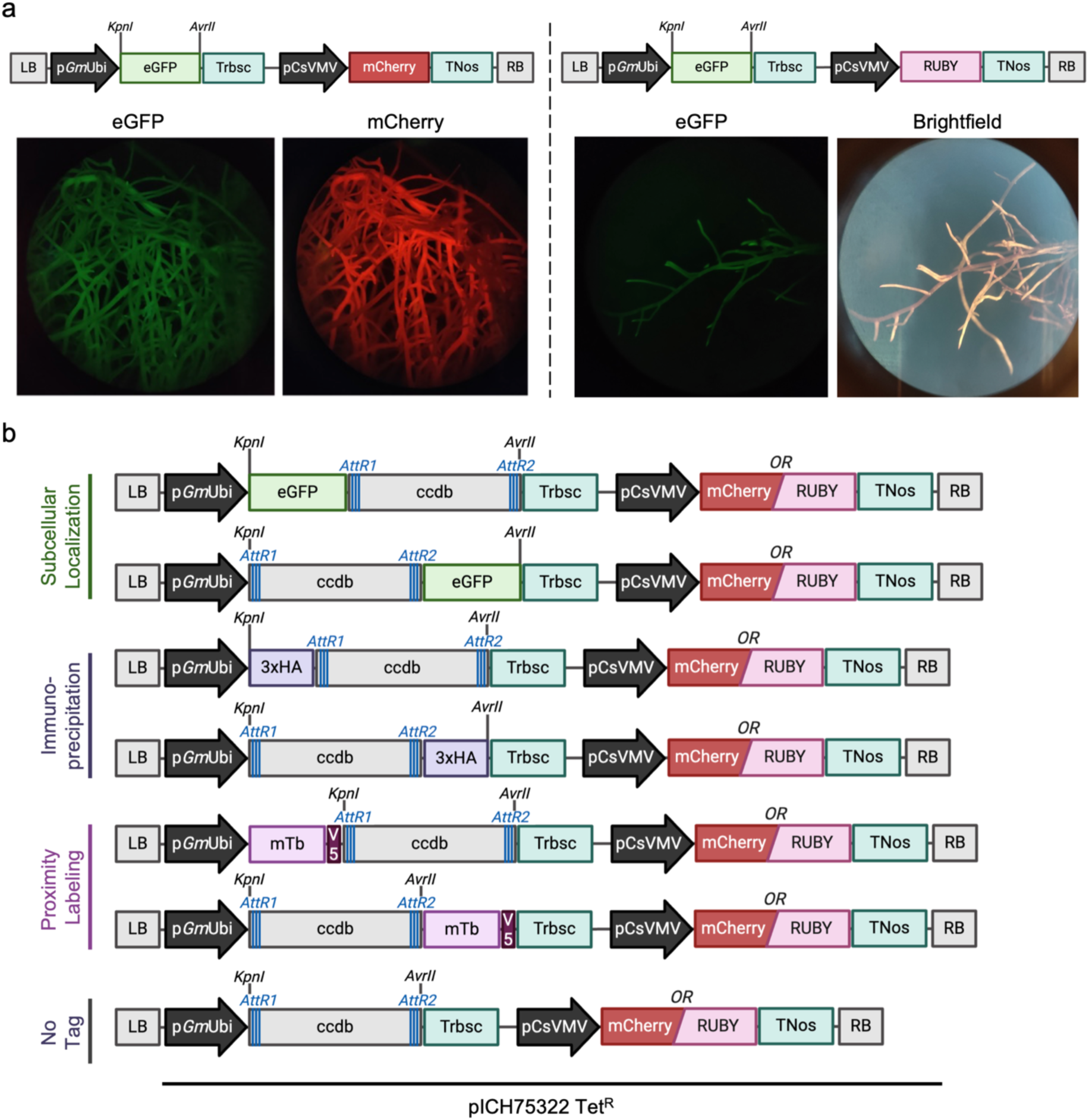
Vectors generated for expression in soybean composite plants. (a) The two intermediate vectors generated prior to GATEWAY conversion. Left: vector expressing GFP and mCherry driven by p*Gm*Ubi and pCsVMV, respectively. Right: vector expressing eGFP and RUBY under p*Gm*Ubi and pCsVMV, respectively. The *KpnI* and *AvrII* sites used for GATEWAY conversion are shown. Root fluorescence was monitored under an epifluorescence binocular using GFP or mCherry filters. RUBY production was monitored using Brightfield. (b) List of the GATEWAY-compatible vectors usable for subcellular localization, immunoprecipitation, proximity-labelling and for overexpression (without a tag). Vectors are available using either mCherry or RUBY as a selectable marker. Created with BioRender.com (agreement number: KN273LHY2A).

During the cloning step into pMiniT2.0, the GFP cassette was flanked by *KpnI* and *AvrII* restriction sites in order to facilitate conversion of these intermediate vectors into GATEWAY-compatible vectors. We replaced the eGFP cassette with different AttR1-ccdB-AttR2 cassettes using restriction-ligation with *KpnI* and *AvrII* (Figure 2b), which were amplified from the pGWB (Nakagawa et al., 2007) and pK7 (Karimi et al., 2002) vector backbones, except for the miniTurbo-ID-V5 cassette which was built independently through an intermediate step. If needed, these two intermediate vectors could be further converted into additional GATEWAY-compatible vectors by replacing the GFP cassette with another GATEWAY cassette such as FLAG-AttR1-ccdB-AttR2, Myc-AttR1-ccdB-AttR2, or V5-AttR1-ccdB-AttR2 using the *KpnI* and *AvrII* restriction sites. To test the functionality of the ccdB selection cassette, we transformed these destination vectors into *E. coli* DH5a and DB3.1 strains and confirmed that only the DB3.1 strain can survive on LB+Tet^5^ (Figure S1). Each of the final destination vectors contained two transcriptional units. The first transcriptional unit contained a recombinant ccdB cassette in frame with either eGFP, 3xHA, or miniTurbo-ID-V5 in both N-terminal and C-terminal fusions, while the second transcriptional contained either the reporter gene mCherry or RUBY (Figure 2b) for a total of 14 destination vectors.

### Expression of subcellular markers in soybean roots

Plant biologists frequently perform subcellular localization studies of GOI in *N. benthamiana* leaves due to the convenience of this model system. Indeed, *N. benthamiana* leaves are suitable for agroinfiltration, and transgenes are expressed in only two to three days. However, the subcellular localization of a protein of interest in *N. benthamiana* leaves may differ compared to soybean roots due to different physiology of the tissue and plant species. With our new set of GATEWAY-compatible vectors, we investigated the potential of using this system to observe subcellular localization of GOIs directly in soybean roots. To this end, we selected and designed various subcellular markers targeting different subcellular compartments, including the nucleoplasm and cytoplasm (eGFP), nucleus (NLS-YFP), microtubules (GFP-MAP4/MBD), actin filaments (LifeActin-mCherry), endoplasmic reticulum (AtWAK2-mCherry-HDEL), plasma membrane (PIP2-GFP), and plasmodesmata (PDLP1-GFP). To test the functionality of our vectors, we shuttled these markers into our set of destination vectors using LR reactions (recombination between AttL1 and AttL2 sites with AttR1 and AttR2 sites using LR clonase II from Invitrogen) to create *in-frame* fusions of the subcellular markers with fluorescent tags.

To rapidly generate transgenic roots, we utilized a modified version of a recently developed protocol, which allows soybean composite plant generation in one-step (see Methods and Fan et al. 2020). Using this system, at least two-thirds of the plants produced transgenic roots after three weeks, making this approach reliable and simple to perform. We observed the expected localization for each subcellular marker in transgenic soybean roots using a LSM700 confocal microscope (Figure 3), highlighting the utility of our vectors for subcellular localization studies in soybean composite plants.

**Figure 3.**
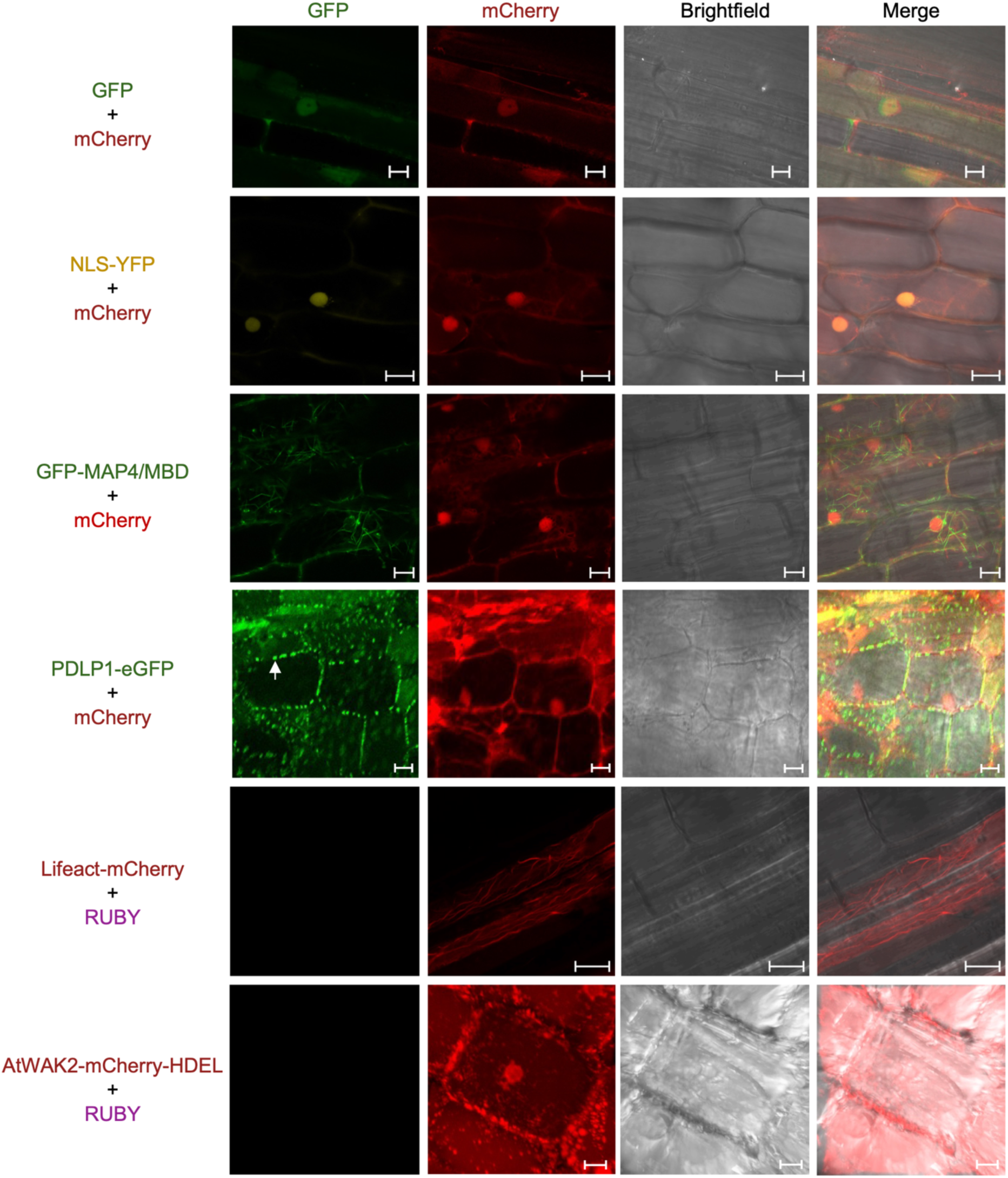
Direct observation of subcellular localization of different subcellular compartment markers. The top line shows the fluorescence channel used (GFP, mCherry, or brightfield). The proteins expressed in soybean composite roots are indiciated on the left. The white arrow indicates plasmodesmata. Scale bar = 10 µm. Experiments were repeated at least twice for each construct.

We next explored the possibility of utilizing the ClearSee clearing method (Kurihara et al. 2015) to achieve 3D reconstruction of these fluorescent signals in soybean composite roots (Supplemental video 1). However, given that soybean roots are thicker than *A. thaliana* roots, an additional step was required in which the roots were manually longitudinally sectioned using a razor blade before fixation and clearing with ClearSee. This step allowed us to obtain high-quality images of the soybean root architecture using Calcofluor White, which stains cell walls, while maintaining the expected subcellular localization of the fluorescent markers (Figure 4, Supplemental video 1). By combining our new set of vectors with the ClearSee method, plant biologists can now rapidly investigate the subcellular localization of a protein-of-interest and monitor any potential impact of a transgene in soybean roots at both the subcellular level and root architecture level.

**Figure 4.**
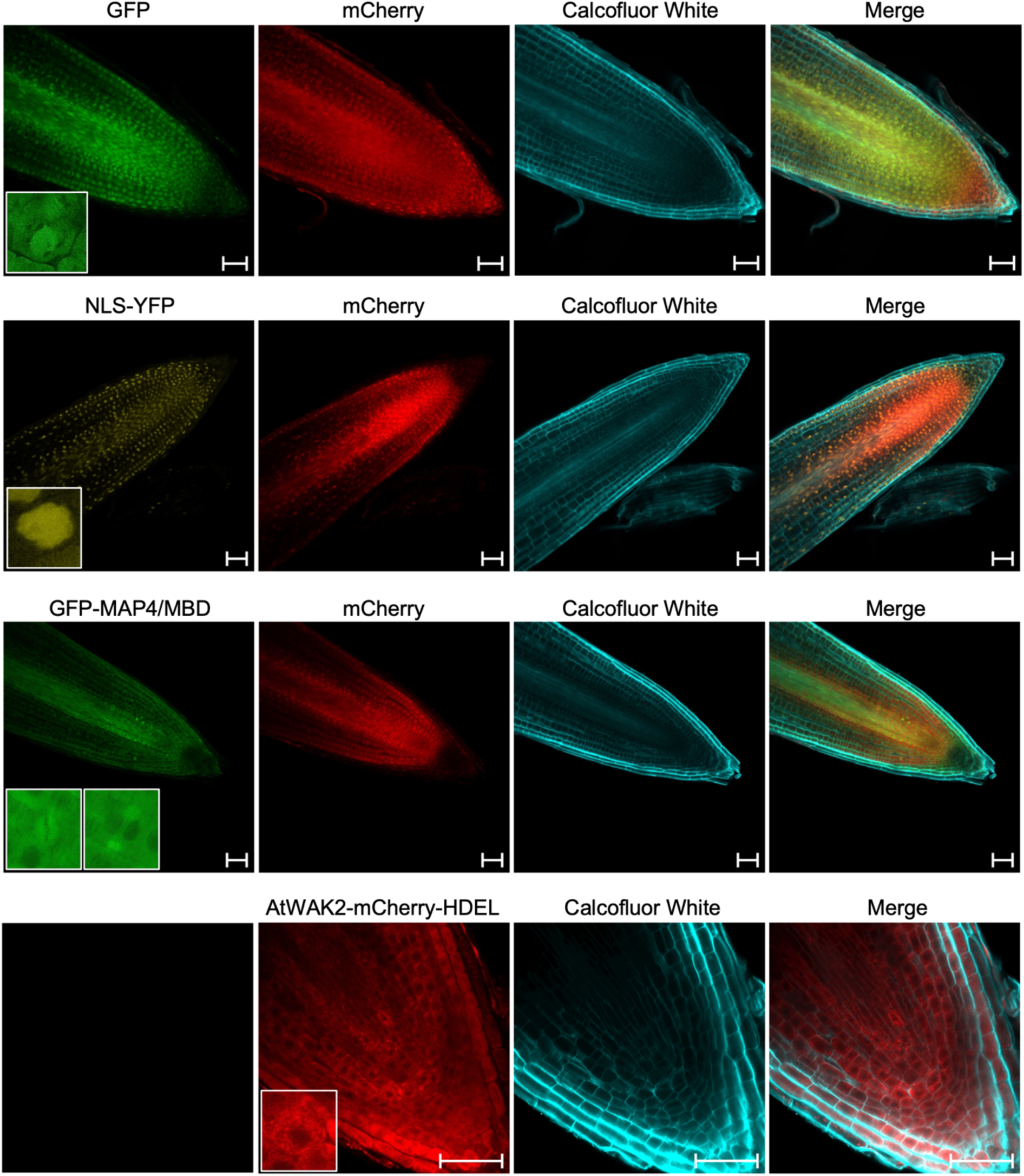
Subcellular localization of four subcellular localization markers in soybean composite roots treated with ClearSee. The fluorescence channel used is indicated across the top of the figure (GFP, mCherry, or Calcofluor white) and the subcellular marker is indicated on the left. Scale bar = 50 µm. Experiments were repeated twice.

Interested in plant-nematode interactions, we were curious if this system could allow us to quickly examine the subcellular localization of a protein-of-interest in the feeding site of plant-parasitic nematodes, such as giant-cells induced by root-knot nematodes (RKNs). To do so, we inoculated soybean composite roots expressing two of the subcellular markers (PDLP1-GFP and PIP2-GFP) with the RKN *Meloidogyne incognita* and the fluorescence of marker proteins under a confocal microscope after hand-cutting the collected RKN-induced galls using a razor blade (Figure 5). As a proof of concept, we successfully observed the expression of the plasma membrane marker (PIP2) as well as the plasmodesmata marker (PDLP1) in giant-cells at 14 dpi (Figure 5). Giant-cells are considered transfer cells that are highly connected to the vascular system, in order to withdraw host nutrients. We observed that the plasmodesmata marker showed high connection between the giant-cells and the neighboring cells as expected. Interestingly, we also observed a strong signal in the nuclei (Figure 5) for this subcellular marker that is not observed in uninfected roots (Figure 2). This could be either due to cleavage of the GFP chimeric protein creating an artifact or could highlight an interesting feature of plasmodesmata regulation in giant-cells. These results show that the vectors can be successfully used to observe subcellular localization in the feeding sites of nematodes.

**Figure 5.**
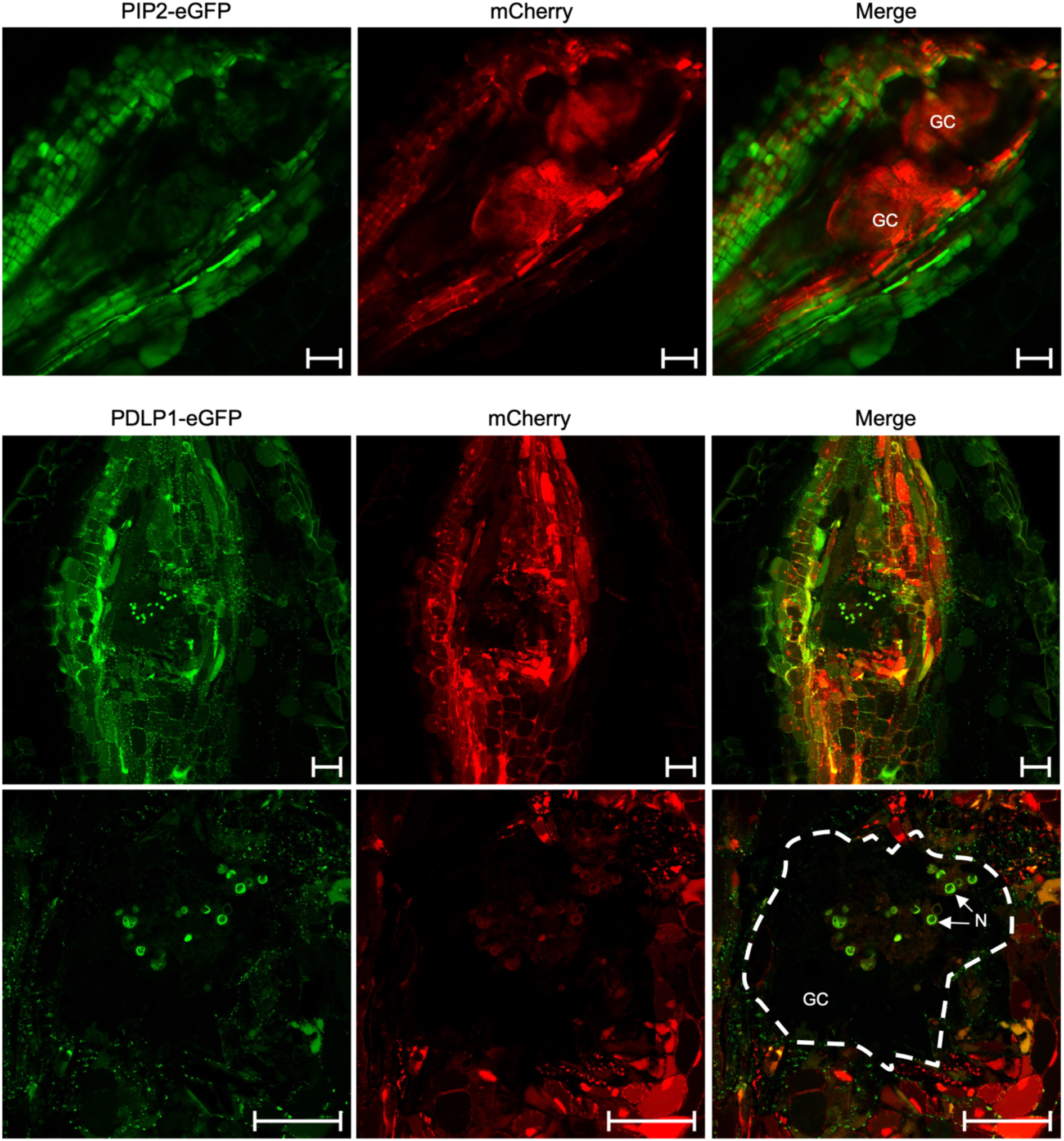
Subcellular localization of PIP2-eGFP and PDLP1-eGFP in giant cells induced by RKN at 14 dpi. GFP fluorescence was used to monitor expression of the PIP2-eGFP (plasma membrane) or PDLP1-eGFP (plasmodesmata) marker genes. The mCherry fluorescence channel was used to monitor the selectable marker for transgenic roots. N = Nucleus; GC = Giant cell; The dotted line contours the giant cell. Scale bar = 50 µm.

### Promoter expression analysis in soybean roots

Since our vectors carry GC-rich restriction sites around the *Gm*Ubi promoter, we investigated the expression strength and localization of several additional promoters which we introduced using restriction ligation in this GC-rich region. We compared the expression of the RUBY reporter gene under the control of the *Gm*Ubi promoter, the *Gm*Actin promoter, P35S+omega (Ω) enhancer viral promoter and the pCsVMV viral promoter (Figure 6a). This allowed us to classify the strength of these promoters from weak to strong in the following order: p*Gm*Actin < P35S + Ω enhancer < pCsVMV < p*Gm*Ubi. All promoters were ubiquitously expressed, except for pCsVMV, which appeared to be less active in the elongation zone of lateral roots. Based on our analysis, the *Gm*Ubi promoter used to drive the expression of the GOI in the vector set is the strongest promoter and is active in every part of the root, making it a good choice for overexpression of GOIs. Additionally, the G*m*Actin promoter can be used to moderately express a GOI.

**Figure 6.**
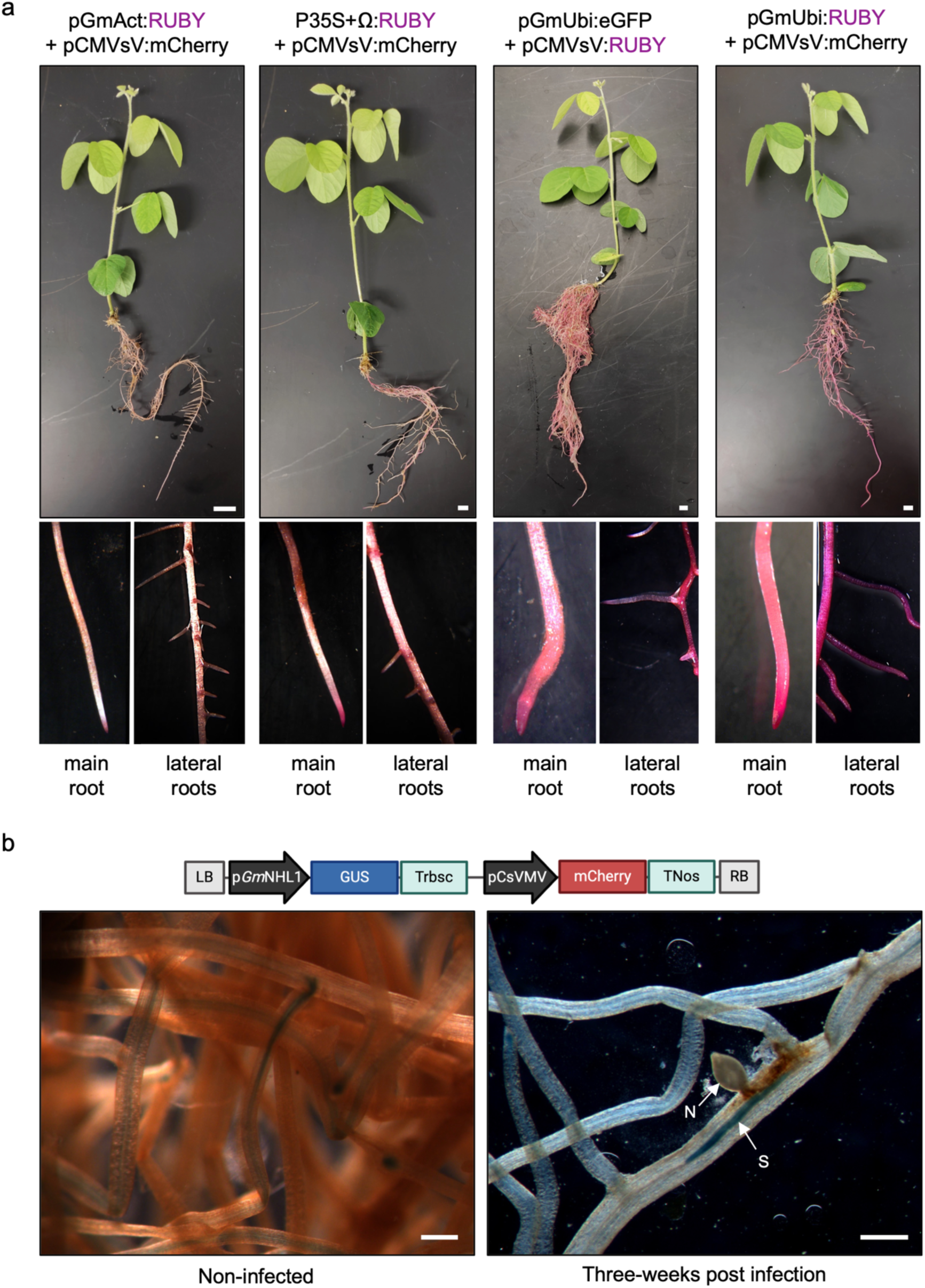
Promoter expression analysis in soybean composite roots. (a) Analysis of RUBY expression in roots driven by four different promoters (indicated above the pictures). Scale bar = 1 cm. Zoomed in pictures of main roots and lateral roots for each construction are shown directly below. (b) GUS expression driven by the *p*NHL1 promoter in uninfected (left) or SCN infected roots (right) three weeks after infection. N = nematode; S= syncytium. Scale bar = 200 µm. Experiments were conducted using 3-4 independent plants. Created with BioRender.com (agreement number: KN273LHY2A).

Cyst nematodes induce the formation of a specialized feeding site known as a syncytium (Davis et al. 2008). We therefore cloned the p*Gm*NHL1 (Glyma03g35930) (Yeckel 2012) to determine if we could express a GOI in soybean composite roots, specifically in syncytia. We cloned 1399 bp of the p*Gm*NHL1 promoter upstream of the ꞵ-glucuronidase (GUS) reporter gene. We observed a clear GUS signal in syncytia of the soybean cyst nematode (SCN) three weeks after infection (Figure 6b). However, we also observed that the p*Gm*NHL1 promoter was expressed in the vascular system of lateral roots in uninfected plants (Figure 6b). Nevertheless, the expression under the p*Gm*NHL1 promoter was clearly enriched in syncytia and, therefore, could be useful for investigating protein function in this structure. Together, our promoter analyses demonstrate how our vectors can be easily and efficiently modified in two cloning steps to replace existing promoters with any number of additional soybean promoters including native promoters, such as p*Gm*NHL1, driving the expression of a fusion protein.

In order for RUBY to be a useful selection marker for gene function analysis during nematode infection, the production of betalain should not have an effect on nematode infection or cyst development. To our knowledge, SCN infection assays using RUBY have not been investigated. For this purpose, we generated soybean composite plants using the two intermediate vectors: p*Gm*Ubi:eGFP + pCsVMV:mCherry and p*Gm*Ubi:eGFP + pCsVMV:RUBY. One month after infection by SCN, we did not observe a significant impact on either the number or size of SCN cysts using RUBY versus mCherry as reporter gene (Figure 7a). Interestingly, during this assay we observed the natural formation of nodules on some composite plants expressing the RUBY reporter gene, indicating that this gene does not seem to prevent nodule formation (Figure 7b), although a more controlled investigation of nodule formation during rhizobial symbiosis in soybean will need to be conducted to validate these observations.

**Figure 7.**
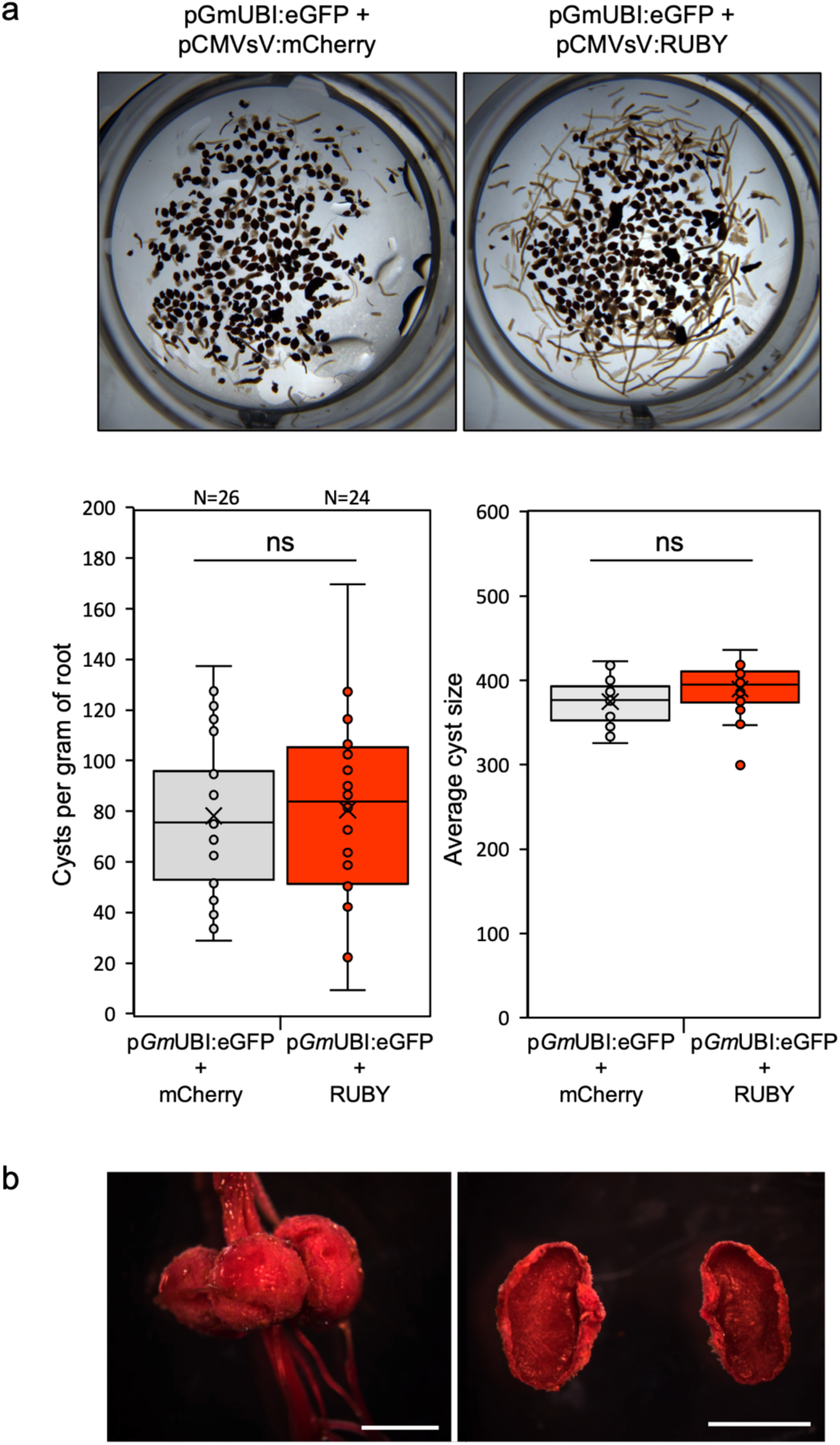
The RUBY reporter gene does not impact SCN infection. (a) Top: representative pictures of the cysts collected from the two constructions harboring the mCherry or RUBY selectable markers. Bottom: quantification of cyst number per gram of root and average size of individual cysts. Experiments were conducted twice. Statistical analysis was conducted by performing a Student’s t-test using p<0.05 as a significance threshold. (b) Picture of natural formation of nodules on soybean composite plant expressing RUBY. Scale bar = 1 cm.

### Expression and immunoprecipitation of a bacterial effector protein

An advantage of using soybean composite roots for gene function analysis, over transient expression in *N. benthamiana*, could be the identification of host proteins that could be targeted by soybean pathogen effectors during infection. In lieu of producing stable transgenics, a large number of proteins could be expressed using this system, allowing high throughput investigation of effector-target interactions. We, therefore, expressed eGFP-3xHA, as well as the AvrPphB^C98S^, a catalytic mutant of a cysteine protease effector from *Pseudomonas syringae*, fused to 3xHA, in soybean composite roots (Figure 8a) and monitored expression by immunoblot analysis. We observed a high level of both chimeric constructs in total protein extracts (Figure 8b). We were able to enrich these proteins by IP (Figure 8b), which could be used to detect effector targets by mass spectrometry.

**Figure 8.**
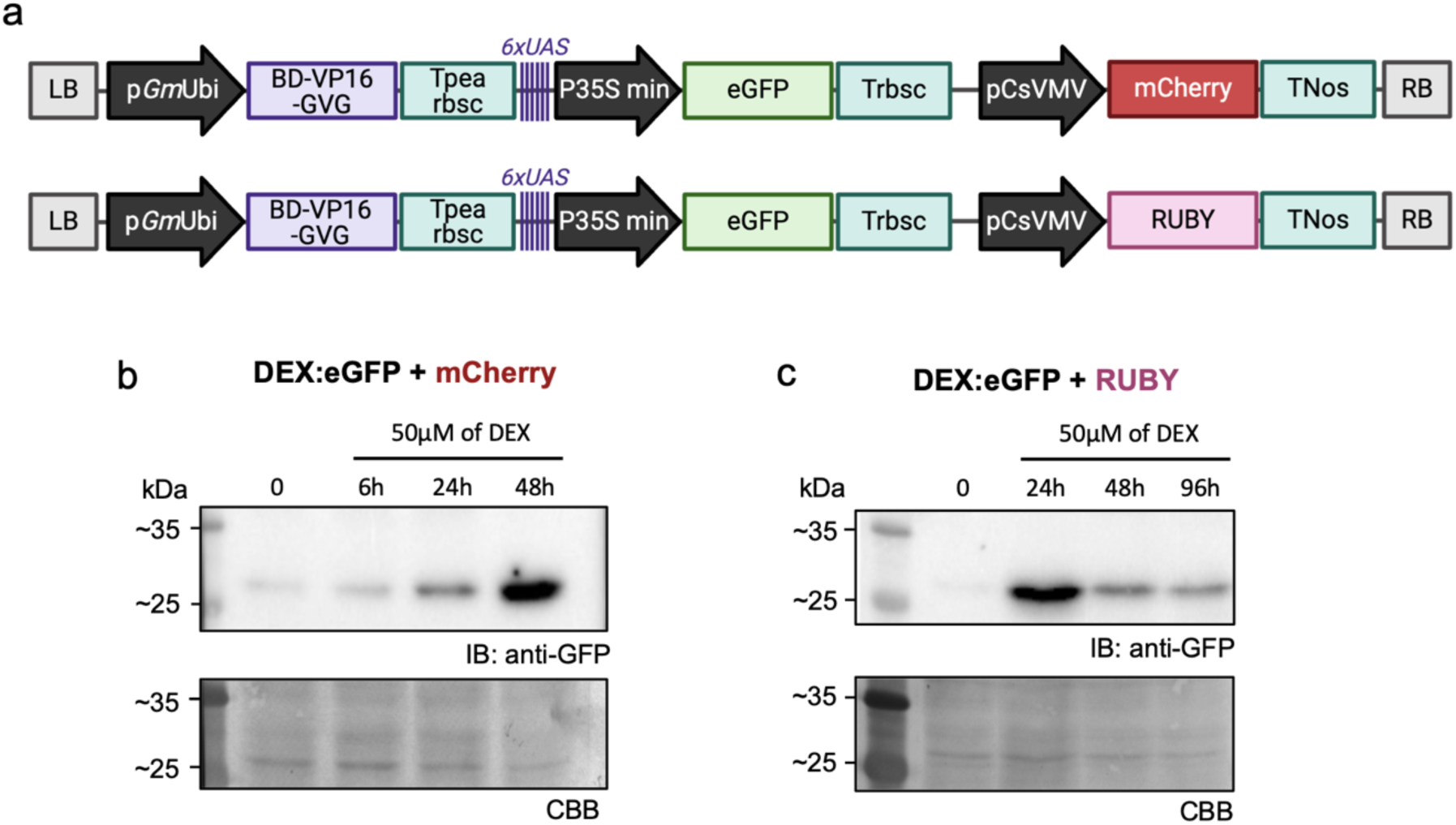
Analysis of DEX-inducible expression in soybean composite roots. (a) Schematic representation of the two intermediate vectors generated before GATEWAY conversion. Immunoblot showing inducible expression of eGFP proteins using (b) mCherry or (c) RUBY as selectable markers. IB = Immunoblot; CBB = Coomassie Brilliant Blue. Created with BioRender.com (agreement number: KN273LHY2A).

In recent years, proximity labeling has become a widely used approach for identifying transient interactions and protein complexes *in planta* (S.-L. Xu et al. 2023). Therefore, we additionally generated GATEWAY vectors that fuse a protein of interest to miniTurbo-ID, a protein that biotinylates proteins in close proximity to the fusion proteins and verified that the full-length fusion protein is efficiently expressed in soybean composite roots (Figure 8c). This iteration of miniTurbo-ID has recently been successfully used to identify a SCN effector target in soybean composite roots (Margets et al. 2024). We therefore expect that these GATEWAY compatible vectors could be highly efficient tools for high throughput identification of effector-host interactions in soybean roots.

### Inducible expression in soybean roots using a dexamethasone-responsive promoter

We generated an additional set of 14 similar vectors as previously described, with a DEX-inducible promoter instead of the constitutive *Gm*Ubi promoter (Figure S2). We selected the DEX-inducible promoter from the pTA7001 vector (Aoyama and Chua 1997) and tested the functionality of these vectors by vacuum infiltrating the roots of soybean composite plants expressing eGFP under this promoter (Figure 9a) with 0.25X Gamborg’s + 50 µm DEX. We observed very little expression in the absence of the inducer in plants expressing mCherry (Figure 9b) or RUBY (Figure 9c) reporter genes. In contrast, a high level of eGFP expression was observed at 48 h and 24 h post-induction, respectively, for plants expressing the mCherry and RUBY reporter genes. Using the vectors generated here, a protein that exhibits toxic or pleiotropic effects on soybean plants can be expressed and used to identify interacting proteins without consequences to the host plant. This system could also be useful for monitoring interactions at specific time-points during the infection process if coupled with proximity labeling or IP fusions.

**Figure 9.**
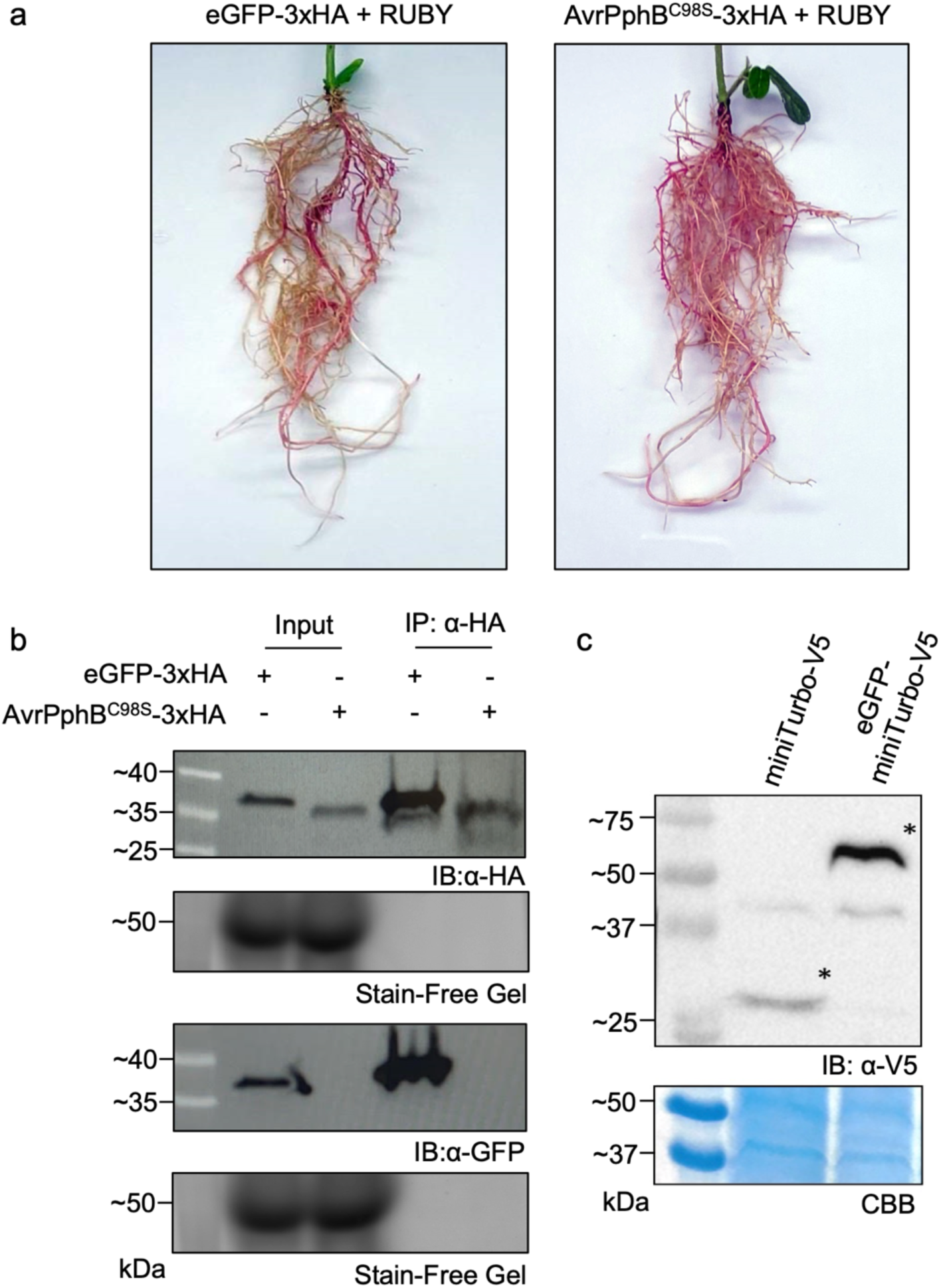
Expression analysis of constructs designed for immunoprecipitation or proximity labeling experiments. (a) Representative picture of the soybean composite plant expressing the RUBY reporter gene as well as eGFP (control) or the catalytically inactive AvrPphB^C98S^ bacterial effector protein. (b) Immunoprecipitation of the AvrPphB^C98S^-3xHA or eGFP-3xHA proteins from soybean composite roots using anti-HA beads. Input lanes correspond to total protein extracts and IP lanes correspond to the protein fraction eluted after IP. Proteins were visualized by chemiluminescence using α-HA or α-GFP primary antibodies and HRP-conjugated secondary antibodies. Stain-free gels were used to assess protein normalization of protein inputs prior to membrane transfer. (c) Immunoblot analysis of miniTurbo-V5 or eGFP-miniTurbo-V5 expression in soybean composite roots. The total protein extract was blotted and visualized using α-V5 primary antibody and an HRP secondary antibody by chemiluminescence. IB = Immunoblot; CBB = Coomassie Brilliant Blue.

## Discussion

Phytopathogens secrete numerous effectors that are intricately regulated throughout the infection process, making it crucial to identify the cellular processes they target to develop novel plant resistance strategies (Mejias et al. 2019; de Almeida Engler et al. 2005; Mitchum 2016). Soybean, an essential crop for both animal and human consumption, faces significant threats from the plant-parasitic nematodes SCN and RKN (Kantor, Eisenback, and Kantor 2024). While transgenic plants are vital for studying effector functions and their targets, generating stable transgenic soybean lines remains labor-intensive and time-consuming, limiting their use for high-throughput screening of large numbers of effectors. Soybean composite plants are now commonly used to assess the importance of given genes in abiotic stress (Hou et al. 2022; Mazarei et al. 2023), SCN (Hawk et al. 2024, 2023; Piya et al. 2022) and RKN virulence (Zhang et al. 2023), as well as interactions with root symbionts (Zhang et al. 2023; Wu et al. 2023). Here, we developed a novel set of GATEWAY vectors to simplify cloning of pathogen genes, such as effectors, or their soybean targets with epitope tags that can be used to investigate pathogen-root interactions in composite soybean plants. We have demonstrated that these vectors can be used to easily generate fusion proteins to confirm protein overexpression, investigate subcellular localization, and generate fusion proteins used for identifying effector-host target interactions through co-IP or proximity labeling.

Our overall goal was to make the assessment of root-pathogen interactions in soybean more efficient and less labor intensive. The use of this new set of vectors, coupled with the generation of soybean composite plants, allows researchers to assess gene function and protein-protein interactions in roots, within two months (Figure 1), from cloning to functional analysis. To do so, we generated a set of GATEWAY destination vectors compatible with the *A. rhizogenes* strain K599 with *in-frame* epitope tags and incorporated a RUBY screenable marker (Figure 2b), eliminating the need for specialized equipment and greatly reducing the labor-intensive nature of selecting transgenic roots. Moreover, this set of vectors can be used in the common heterologous system such as *N. benthamiana* or *A. thaliana* since they can also replicate in *A. tumefaciens*. We showed that the RUBY reporter gene did not impact SCN or potentially Rhizobial infection (Figure 7a), highlighting its usefulness in transgenic root selection and the investigation of root-microbe interactions. Thus, this system makes it possible to conduct high throughput screens for effector function and the identification of host-interacting proteins, which has been a major bottleneck for studying plant-microbe interactions in soybean.

Very few soybean pathogen effectors have been functionally characterized, in part because of the labor-intensive nature of generating transgenic plants for protein expression and analysis in soybean. Subcellular localization of effectors, and their host target proteins, is essential for providing clues about the function of pathogen effectors. We have shown that, coupled with or without ClearSee, the subcellular localization of a given protein in soybean roots (Figures 3 and 4), or in nematode feeding sites (Figure 5), is now easily achievable. Indeed, almost all studies on nematode effectors use heterologous systems such as onion epidermis or *N. benthamiana* leaves to decipher the subcellular localization of a given effector and therefore cannot be investigated directly within the nematode feeding sites. In *N. benthamiana* or onion epidermis the expression of nematode effectors has been shown to alter the localization of host proteins targeted by the effectors (Pogorelko et al. 2019; Hewezi et al. 2015). Moreover, nematode feeding sites cause a dramatic reorganization of plant cell architecture which could also affect host protein localization (Favery et al. 2016; Ohtsu et al. 2017; Gheysen and Mitchum 2011). We therefore see high utility in the use of this system for investigating the localization of both effector proteins and host-interacting proteins in giant-cells or syncytia produced during nematode infection.

We additionally generated a set of vectors that express GOIs under DEX-inducible promoters (Figure S2) that show little to no protein expression in the absence of DEX (Figure 8b-c). These constructs will be useful for expression of toxic proteins (Verma et al. 2018; Mejias et al. 2021; Vijayapalani et al. 2018) or proteins that drastically alter root development (Vieira and Gleason 2019), which can impede downstream analysis. For example, the avirulent AvrPphB effector leads to a hypersensitive cell death response (Zhu et al. 2004) when expressed in plant expressing the corresponding resistance gene, making it impossible to investigate interactions in soybean using the catalytically active variant of this protease. Achieving precise temporal expression of effectors will also aid the identification of host-interacting proteins that occur at specific points of infection. Ectopic expression of effectors that are expressed constitutively throughout infection can cause artifacts that do not necessarily reflect their role in pathogen virulence. Therefore, the addition of a set of inducible promoters will also strengthen the identification of true host-protein interactions. By doing so, we expect this system could be used to effectively identify host susceptibility genes that are hijacked by effectors to promote infection. The modification of susceptibility genes to evade effector detection is currently one of the most effective and sustainable approaches for improving resistance against plant pathogens (Wang et al. 2024). We have also modified these vectors so that pathogen or host genes can be expressed under their native promoter, by replacing the *Gm*Ubi promoter through digestion-ligation cloning. This will also allow researchers to study the effects of protein expression on host immunity or development under more “natural” conditions.

The flexibility of these vectors, and easy replacement of the *Gm*Ubi promoter, also open possibilities to modify the vectors for use in other crop plants that are not easily transformed. For example, these vectors could be used to generate composite plants such as tomato, cucumber or even woody plants such as prunus species (Ho-Plágaro et al. 2018; Y. Fan et al. 2020; Bosselut et al. 2011). Here, we demonstrate the possibility of these vectors for studying plant-nematode interactions, and expression of a catalytically inactive variant of the avirulent bacterial effector protein (AvrPphB) from *Pseudomonas syringae* (Zhu et al. 2004). However, soybeans, like most crops, are susceptible to a number of additional root pathogens of economic importance that currently lack understanding of molecular host-microbe interactions including filamentous pathogens such as fungi (for example, *Fusarium* species) and oomycetes (for example, *Phytophtora sojae* or *Pythium* species). Therefore, these vectors can be used to investigate pathogen-root interactions in numerous pathosystems with very little modification.

A limitation of the composite plant system is that the transgenic plants are not stable. Recently, a grafting method using Arabidopsis-overexpressing Cas9 line alongside a grafted root system expressing a guide RNA (gRNA) fused to a mobile signal allows the gRNA to reach reproductive organs (Yang et al. 2023). This system could provide the ability to bypass the regeneration steps needed to produce stable transgenic plants. It would be exciting to test in the future if transgenic roots generated from soybean Cas9 expressing lines could serve as a source to produce mobile gRNA that would reach the reproductive aerial organs and therefore bypass regeneration to allow precise heritable gene edits.

In conclusion, we have developed a novel set of GATEWAY-compatible vectors that can be used to investigate cell biology studies such as host-microbe interactions in soybean composite roots. The vectors are designed for flexible modification of promoters driving GOI expression and the incorporation of new epitope tags, and allow easy identification using mCherry or the RUBY reporter genes. We envision that the utility of these vectors can be extended to other composite plant systems allowing *in planta* investigation of pathogen effectors in their host systems, rather than relying on the use of heterologous systems such as *N. benthamiana*.

## Data Availability

All vectors generated during this study are available upon request from the corresponding author at Iowa State University.

## Acknowledgements

The pBR322 vector used for Tetracycline cassette amplification was kindly provided by Dr. Gwyn Beattie (Iowa State University). pGWB414 and pGWB415 was a gift from Tsuyoshi Nakagawa (Addgene plasmid # 74808, http://n2t.net/addgene:74808, RRID:Addgene_74808 for pGWB414; and Addgene plasmid # 74809, http://n2t.net/addgene:74809, RRID:Addgene_74809 for pGWB415). pICH75322 was a gift from Sylvestre Marillonnet (Addgene plasmid # 48051, http://n2t.net/addgene:48051, RRID:Addgene_48051). The pK7WGF2, pK7GWF2 and pK2GW7 was a gift from Mansour Karimi and VIB (Ghent) collection (https://vectorvault.vib.be/collection/). pENTR221-mCherry was a gift from Peter Jon Nelson (Addgene plasmid # 79509, http://n2t.net/addgene:79509, RRID:Addgene_79509). 35S:RUBY was a gift from Yunde Zhao (Addgene plasmid # 160908, http://n2t.net/addgene:160908, RRID:Addgene_160908).

## Author Information

JM and TJB designed the experiments. JM, AM, MB, JF, EK, JG and TRM conducted the experiments. JM and MB wrote the manuscript with input from all authors. TJB, RWI, and SAW supervised and secured funding.

## Ethics Declarations

The authors declare no competing interests.

