## Supplemental Data for "A novel toolbox of GATEWAY-compatible vectors for rapid functional gene analysis in soybean composite plants"

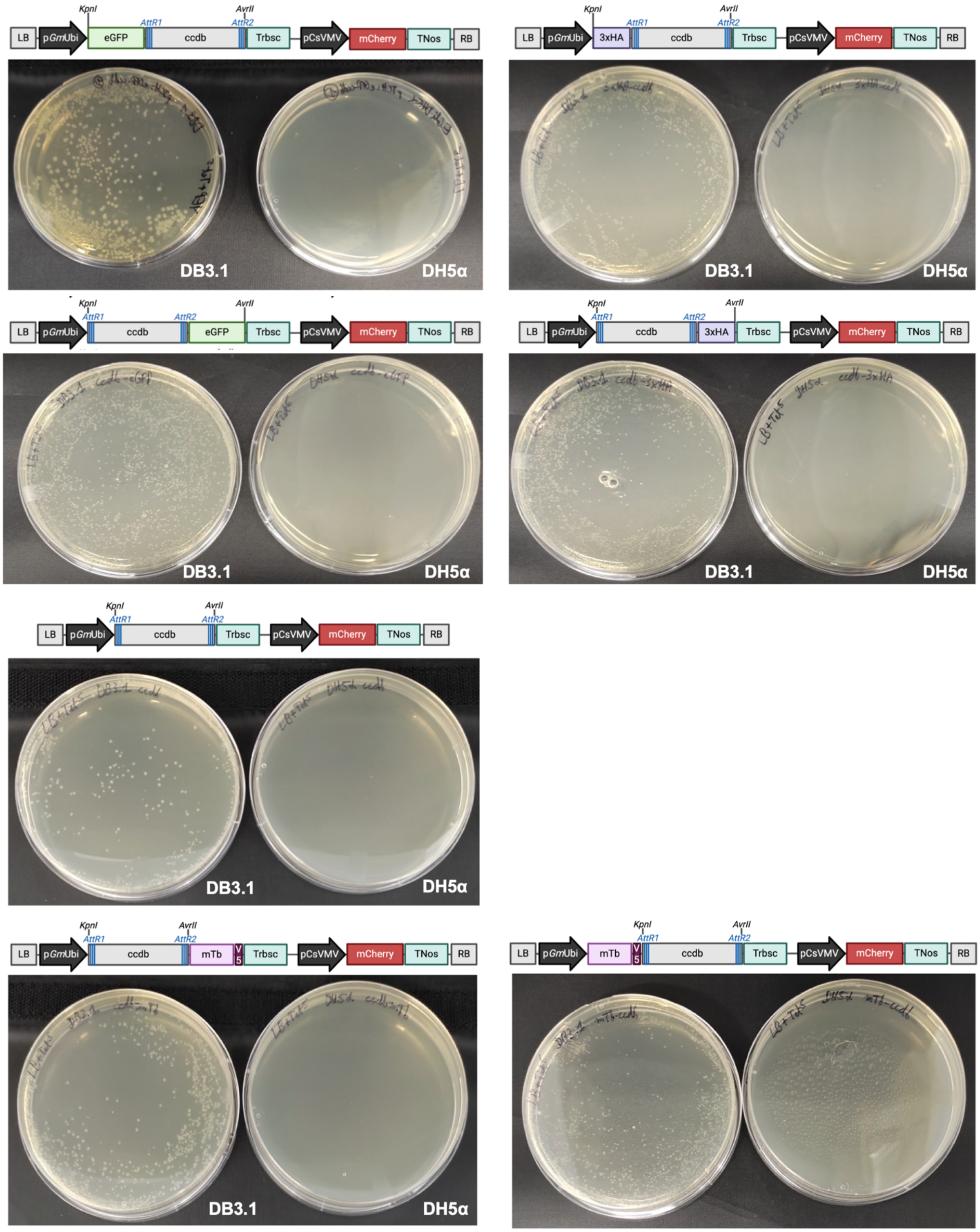


**Figure S1.** Efficiency of ccdb counter selection in the GATEWAY vector set. 50 ng of the vectors was transformed into *E. coli* DH5α or DB3.1. Created with BioRender.com (agreement number: RS273LI22Z).

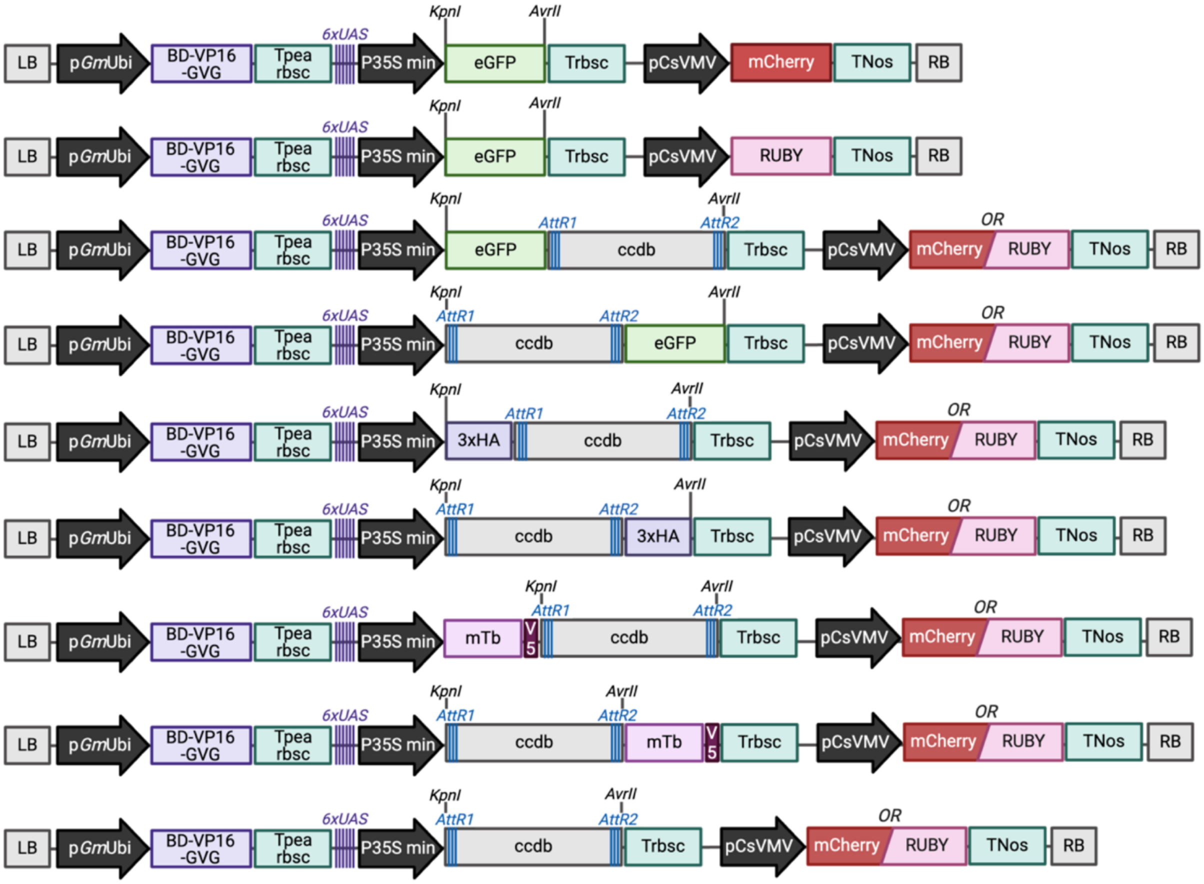


**Figure S2.** Schematic representation of all the T-DNA regions of the set of dexamethasone-inducible vectors available for soybean composite plant generation. Created with BioRender.com (agreement number: QF273LIAYS).

**Table S1**. List of primers used in this study.

| Purpose | Units GG | Primers name | Sequence (5'-3') | GG overhang |
|---|---|---|---|---|
| *Replacing KanaR by TetR* | | **SspI_Tetc-F** | GCAATATTttcaagaattctcatgtttgacag | |
| | | **SspI_Tetc-R** | CGAATATTatgcgcacccgtggccaggaccc | |
| *GoldenGate units* | *Unit 1* | **PGmUBI1_GG_F1** | tgccCCGCGGTCCGGAgtaacagataaagcgaagcttg | TGCC-**Unit1**-CTTA |
| | | **PGmUBI1_GG_F2** | ggctacggtctcc***tgcc***CCGCGGTCCGGAgtaacagataaag | |
| | | **PGmUBI1_GG_R1** | GGCGCGCCCCCGGGggtgcggccgctgtcgagtcaac | |
| | | **PGmUBI1_GG_R2** | ggctacggtctca**taag**GGCGCGCCCCCGGGggtg | |
| | *Unit 2* | **eGFP_GG_F1** | GGTACCacaaaaATGgtgagcaagggcgaggagCTG | CTTA-**Unit2**-TTCG |
| | | **eGFP_GG_F2** | ggctacggtctca***CTTA***GGTACCacaaaaATGgtgaag | |
| | | **eGFP_GG_R1** | CCTAGGTCActtgtacagctcgtccatgccg | |
| | | **eGFP_GG_R2** | ggctacggtctca**cgaa**CCTAGGTCActtgtac | |
| | *Unit2* | **Ruby_middle_GG_F** | ggctacggtctca***CTTA***GGTACCATGGATCATGCGACCCTCGCCATG | |
| | | **Ruby_middle_GG_R** | ggctacggtctca**cgaa**CCTAGGTCACTATCACTGGAGGCTTGGCTC | |
| | *Unit2* | **MCS_mTb_GG_F1** | ACCCCTAGGcggaggtggttctGCAAGGGACCCCCCAGTCGCGAC | |
| | | **MCS_mTb_GG_F2** | ggctacggtctca***CTTA***GGTACCCCTAGGcggaggtggttctGC | |
| | | **MCS_mTb_GG_R** | ggctacggtctca**cgaa**TCAGGTACTGTCAAGACCGAGAAG | |
| | *Unit2* | **mTb_MCS_GG _F** | ggctacggtctca**CTTA**acaaaaATGGCAAGGGACCCCCCAGTCGC | |
| | | **mTb_MCS_GG_R1** | CTAGGGGTACCagaaccacctccGGTACTGTCAAGACCGAGAAGG | |
| | | **mTb_MCS_GG_R2** | ggctacggtctca**cgaa**TCACCTAGGGGTACCagaaccacctcc | |
| | *Unit3* | **TRbsc_GG_F** | ggctacggtctca***TTCG*****G**ttcgagtattatggcattggg | TTCG-**Unit3**-AAGG |
| | | **TRbsc_GG_R** | ggctacggtctca**cctt**gatttgacacatttttactc | |
| | *Unit4* | **PCmVsV_GG_F** | ggctacggtctca**AAGG**ttccagaaggtaattatccaag | AAGG-**Unit4**-GCAA |
| | | **PCmVsV_GG_R** | ggctacggtctca**ttgc**ggatcacaaacttacaaatttctc | |
| | *Unit5* | **mCherry_GG_F** | ggctacggtctca***GCAA***acaaaaATGGTGAGCAAGGGCGAGGAGGA | GCAA-**Unit5**-GATG |
| | | **mCherry_GG_R** | ggctacggtctcc**catc**TCACTTGTACAGCTCGTCCATG | |
| | *Unit5* | **Ruby_GG_end_F** | ggctacggtctca***GCAA***ATGGATCATGCGACCCTCGCCATG | |
| | | **Ruby_GG_end_R** | ggctacggtctcc**catc**TCACTATCACTGGAGGCTTGGCTC | |
| | *Unit6* | **TNos_GG_F** | ggctacggtctcc***gatg***tgcagatcgttcaaacatttggc | GATG-**Unit6**-GGGA |
| | | **TNos_GG_R** | ggctacggtctcct**ccct**catgtttgacagcttatcatc | |
| *Amplifying ccdb cassette* | | **KpnI-eGFP-F** | caGGTACCatggtgagcaagggcgaggagctg | |
| | | **ccdb-AvrII-R** | gtCCTAGGaccactttgtacaagaaagctgaac | |
| | | **KpnI-ccdb-F** | caGGTACCacaagtttgtacaaaaaagctgaac | |
| | | **eGFP-AvrII-R** | gtCCTAGGttacttgtacagctcgtccatgcc | |
| | | **3xHA-KpnI-F** | caGGTACCatgagcgggttaattaacatcttttac | |
| | | **AvrII-3xHA-R** | gtCCTAGGctaagcgctgcactgagcagcg | |
| *pDONR221* | | **NLS-YFP_GW5_ATG_F** | **AAAAAGCAGGCTTC**ACCatgaagcgtcctgctgctactaag | |
| | | **NLS-YFP_GW3_STOP_R** | **AGAAAGCTGGGTG**ttacttgtacagctcgtccatgcc | |
| | | **LifeAct_GW5_ATG_F** | **AAAAAGCAGGCTTC**ACCAtgggtgtcgcagatttgatcaag | |
| | | **LifeAct_GW3_STOP_R** | **AGAAAGCTGGGTG**tcacttgtacagctcgtccatgcc | |
| | | **MAP4MBD_GW5_ATG_F** | **AAAAAGCAGGCTTC**ACCtcccggcaagaagaagcaaaggc | |
| | | **MAP4MBD_GW3_STOP_R** | **AGAAAGCTGGGTG**ttaggcacctcctgcaggaaagtgg | |
| | | **AvrpPhBcat_GW5_ATG_F** | **AAAAAGCAGGCTTC**ACCatggggtgtgcatcctcttcaggc | |
| | | **AvrpPhBcat_GW3_noSTOP_R** | **AGAAAGCTGGGTG**cgaaactctaaactcgtttacgc | |
| | | **GUSplus-GW5-F** | **AAAAAGCAGGCTTC**ACCATGgtagatctgagggtaaatttc | |
| | | **GUSplus-GW3-R** | **AGAAAGCTGGGTG**TCACACGTGATGGTGATGGTGATGgc | |
| | | **AttB1 adapter** | GGGGACAAGTTTGTACAAAAAAGCAGGCT | |
| | | **AttB2 adapter** | GGGGACCACTTTGTACAAGAAAGCTGGGT | |
| Cloning Promoters | | **ApaI_P35S-F** | CAGGGCCCcaatcccacaaaaatctgagcttaac | |
| | | **P35S-AscI-R** | GTGGCGCGCCTGTAATTGTAAATGTAATTGTAATG | |
| | | **ApaI_PromActin-F** | CAGGGCCCCCATGGACTTTAACAGCAACA | |
| | | **PromActin-AscI-R** | GTGGCGCGCCGGTTGTTTAAGGTAAAAGATG | |
| | | **ApaI_PromAtSCR_F** | CAGGGCCCAAGTCTAAAAGGGCAGAAAAG | |
| | | **PromAtSCR_AscI_R** | GTGGCGCGCCGGAGATTGAAGGGTTGTTGGTC | |
| | | **ApaI_PromGmNHL1_F** | CAGGGCCCCAACGGTAAATGCGCCTCCTTCG | |
| | | **PromGmNHL1_AscI_R** | GTGGCGCGCCGCTTTAGGATTGGTGATATAATAG | |
